# A reliable and unbiased human protein network with the disparity filter

**DOI:** 10.1101/207761

**Authors:** Gregorio Alanis-Lobato, Miguel A. Andrade-Navarro

## Abstract

The living cell operates thanks to an intricate network of protein interactions. Proteins activate, transport, degrade, stabilise and participate in the production of other proteins. As a result, a reliable and systematically generated protein wiring diagram is crucial for a deeper understanding of cellular functions. Unfortunately, current human protein networks are noisy and incomplete. Also, they suffer from both study and technical biases: heavily studied proteins (e.g. those of pharmaceutical interest) are known to be involved in more interactions than proteins described in only a few publications. Here, we use the experimental evidence supporting the interaction between proteins, in conjunction with the so-called disparity filter, to construct a reliable and unbiased proteome-scale human interactome. The application of a global filter, i.e. only considering interactions with multiple pieces of evidence, would result in an excessively pruned network. In contrast, the disparity filter preserves interactions supported by a statistically significant number of studies and does not overlook small-scale protein associations. The resulting disparity-filtered protein network covers 67% of the human proteome and retains most of the network’s weight and connectivity properties.

## Introduction

Charting reference maps of protein interactions that take place in the human cell, coupled with the use of network analytics on the resulting human protein network (hPIN), has found many applications. Some examples are protein function determination^1,2^, identification of disease-associated proteins^3–5^ and drug repurposing^6^. These kinds of studies have shed light on the relationship between network structure and complex biological phenotypes and underline the key role that protein interactomics is playing on projects related to human health^7–10^.

Although we are still far from a complete and stable reference hPIN^11–13^, many efforts strive for the systematic mapping of a proteome-scale wiring diagram of the cell^13–17^. However, currently available protein interaction maps have small overlaps between them^13^ and contain high false positive and negative rates^11^. To alleviate these problems and facilitate the construction of reliable subnetworks of the hPIN, protein-protein interaction (PPI) databases, like STRING^18,19^and HIPPIE^20,21^, have established confidence scoring systems for PPIs. STRING’s confidence score corresponds to the probability of finding linked proteins within the same pathway^18^. On the other hand, HIPPIE’s score relies on the amount and quality of the experimental evidence supporting each PPI^20^. The problem is that the construction of high-quality hPINs, only considering interactions with high confidence scores (referred to as global filtering), results in excessively pruned networks and removes potentially interesting information present only below the chosen cutoff^22^.

Another important issue with present-day hPINs is their study bias. Despite several studies describing that disease or essential proteins possess a high number of interaction partners (i.e. they have a high degree)^23–28^, it is becoming evident that this may be due to the number of times that they are used as baits in PPI screenings. In 2011, Brito and Andrews noticed that membrane proteins are underrepresented in PPI datasets when compared to nuclear ones^29^. The reason can be technical (e.g. membrane proteins have low abundances and are thus difficult to detect) but also owed to a bias in favour of nuclear proteins (proteins that translocate to the nucleus are commonly used as drug targets^30,31^). Rolland and colleagues found a strong correlation between the number of studies associated with human proteins and the size of their neighbourhood in the hPIN^15^. Schaefer *et al.* observed the same trend, in addition to a statistically significant difference between the degree of cancer-related proteins and non-cancer ones^32^. However, this difference was not as strong when they compared cancer proteins with non-cancer gene products associated with the same number of studies ^32^. More recently, Luck and colleagues surveyed hPINs built with literature data or functional relationships between proteins and detected the same biases^13^. Using all the experimentally-supported PPIs provided by most molecular interaction databases, we were able to reproduce all these observations (see Fig. 1 and Methods).

**Figure 1.**
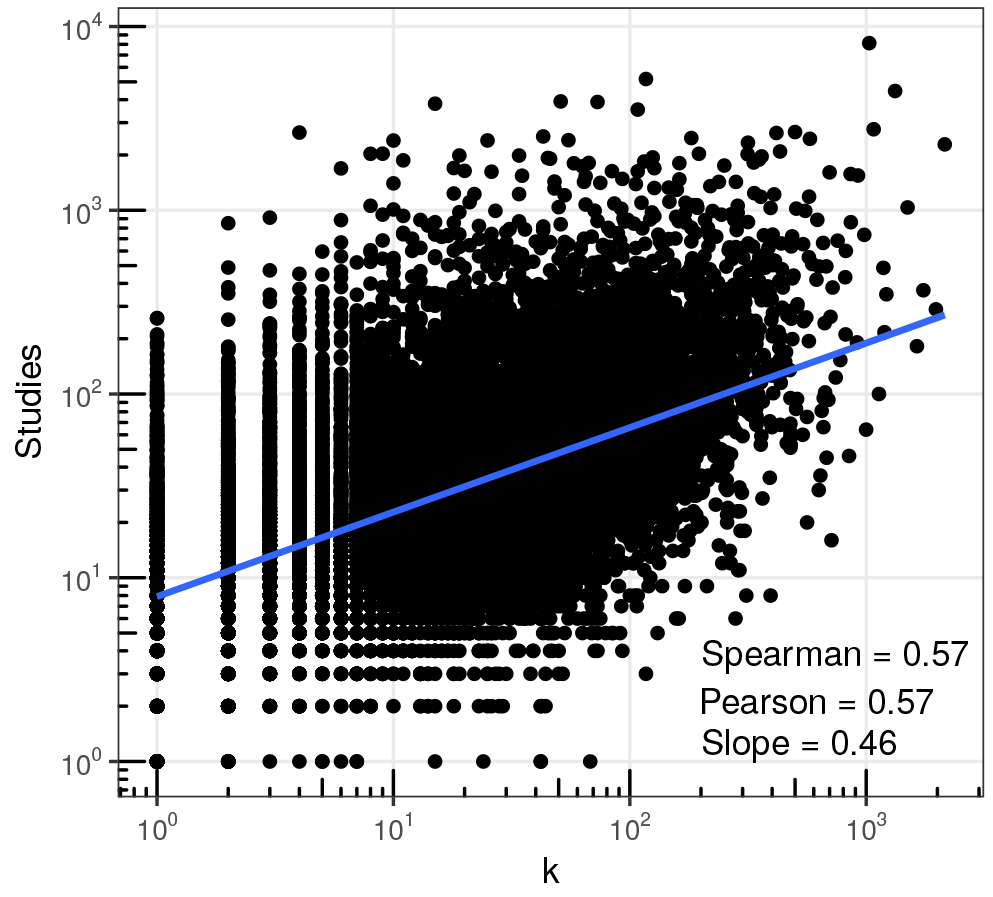
Study bias in the hPIN. The number of studies associated with proteins in the hPIN as a function of their degree *k*. Pearson and Spearman correlation coefficients between the two dimensions are reported, as well as the slope of the blue regression line (see Methods for more details).

In summary, one must carefully consider the incompleteness and biases of the hPIN when drawing conclusions and posing hypotheses based on this data. In this paper, we explore the possibility of constructing a reliable and unbiased human protein interactome, while keeping most of its network components. To that end, we resort to the so-called disparity filter^33^, a tool from the field of network science. This method was designed to extract the most salient connections between the components of a weighted network (i.e. the representation of a complex system where links carry weights denoting their importance) without down-playing small-weight interactions that represent relevant signals at the small scales^33^. Network scientists have used the disparity filter to identify significant trade channels in the international trade system^34^ and to remove links that are weakly related to the overall function of a variety of complex networks^22^.

Our present-day view of the hPIN is incomplete and noisy^11–13^. Pruning this structure without considering its multi-scale nature could result in poor representations of the protein network in the human cell. Consequently, we propose the use of a more educated interaction removal approach that exploits connectivity information at the protein level^33^ and can produce a more reliable and unbiased hPIN, with better coverage.

## Results

### The disparity filter

Consider an undirected weighted network *G* = (*V,E*) with *N*_*tot*_ = |*V*| nodes and *L*_*tot*_ = |*E*| edges. Each edge connects a pair of nodes *i* and *j* and has an associated weight *w*_*ij*_ ∈ (0, ∞) reflecting its importance. If we define the strength of node *i* as *s*_*i*_ = ∑_*j*_ *w*_*ij*_^35^, then the normalised weight of the edges linking *i* with its neighbours is *ω*_*ij*_ = *w*_*ij*_*/s*_*i*_ with ∑_*j*_ *ω*_*ij*_ = 1^33^. Note that this normalisation happens at the level of each node. Thus, *ω_ij_* can be different from *ω*_*ji*_. The disparity filter identifies relevant edges for a node *i* with *k* neighbours by determining the probability that their normalised weights are the result of a random assignment from a uniform distribution^33^ (see Methods). Given this null model, the probability of observing a particular normalised weight *ℓ*, touching a node with degree *k* is given by:

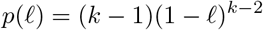

Salient edges are thus those whose normalised weight satisfies the relation:

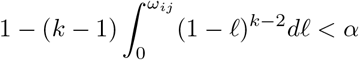

In other words, relevant edges have weights that are statistically greater than what is expected by chance, at the significance level α^33^ (see Methods). Since *ω*_*ij*_ can be different from *ω*_*ji*_, weights can be significant for node *i* but not for *j* and vice versa. The disparity filter keeps edge *ij* if it turns out to be relevant to either node (see Fig. 2). This means that one always considers the minimum value of α related to a particular edge. It is also worth noting that nodes of degree 1 should be treated separately since their only edge always has a normalised weight of 1. In this paper, we keep these edges only if they are significant to the node at the other end. Finally, nodes with degree 0 after filtering are removed from the network. The resulting filtered network *G*_*bb*_, containing the most relevant edges in the system, is hereafter referred to as the backbone of the original one and it has *N*_*bb*_ ≈ *N*_*tot*_ nodes and *L*_*bb*_ ≪ *L*_*tot*_ edges.

Fig. 2 depicts the application of this approach to a weighted toy network and puts it in contrast with global filtering. Guided by cut-points like the upper quartile (400) or the median (16) of the weight distribution to remove edges, global filtering results in node loss and lack of the weight and degree heterogeneities observed in the original network. On the other hand, the disparity filter judges edge importance independently for each node, producing a backbone *G*_*bb*_ with a better resemblance to *G*.

### Application of the disparity filter to the hPIN

We constructed an hPIN with experimentally-supported interactions reported by interaction providers in PSIC-QUIC^36^ (see Methods). The network contains *L*_*tot*_ = 326,758 weighted interactions between *N*_*tot*_ = 16,627 proteins. Edge weights correspond to the number of studies supporting each protein interaction (see Methods, Supplementary Data S1 and S2). The pieces of evidence associated with an interaction are a proxy for reliability. A recent study showed that PPIs reported in two or more publications can be validated at significantly higher rates than PPIs with only one associated study^15^.

Fig. 3 readily highlights the benefits of the disparity filter when contrasted with a global filter, which discards protein interactions with less than *w*_*c*_ associated studies (see Fig. 2). In Fig. 3a, we can see how the resulting fraction of nodes (*N*_*bb*_/*N*_*tot*_) and weight (*W*_*bb*_/*W*_*tot*_) in the backbone remains high, even if we consider very stringent values of α. In contrast, these fractions decrease rapidly in the global filter, and we lose more than 75% of the nodes, even with the less conservative values of *w*_*c*_. The average clustering coefficient of the backbone oscillates around the original value in both scenarios, but we have to consider that in the global filter these values come from small subnetwork remnants of the original hPIN. It is also worth noting that the plots in both panels of Fig. 3a consist of the same number of points. In the global filter, however, these points accumulate to the right side of the plot, where the fraction of remaining links in the backbone (*L*_*bb*_/*L*_*tot*_) is close to zero.

Fig. 3b shows how the cumulative degree distribution of the hPIN does not considerably change when filtered with different values of α. In the case of the global filter, the distribution changes drastically even when we remove edges with low weights. The same occurs with the cumulative weight distribution, as depicted in Fig. 3c: with the disparity filter, the probability of observing small weights does not change as radically (left panel) as with the global filter (right panel).

We also studied how the number of connected components (*CC*) and the size of the largest one (*LCC*) change as increasingly stringent filters are applied to the hPIN. We found that the disparity-filtered hPIN remains connected as a single component up to α ≈ 0.5 (Fig. 4a), whereas a global filter that removes interactions supported by less than 2 studies (*w*_*c*_ = 2). results in an hPIN with only 32% of the input proteins in the largest connected component (Fig. 4b). Fig. 4 also demonstrates that while the disparity filter breaks the hPIN into one component per protein at the extreme case of α = 0, for the global filter this starts to happen when interactions supported by less than 10 studies are removed (*w*_*c*_ = 10). These results allow us to see that the disparity filter retains interactions with both low and high weights, as long as they are important for the proteins at either of their ends. This strategy leads to a filtered network with as many proteins as possible, most of them in the network’s largest connected component, and with a very high fraction of the total input weight.

### Significant protein interactions are highly reliable

To explore whether significant edges, according to the disparity filter, correspond to reliable protein interactions, we monitored the fraction of PPIs in hPINs pruned with the global filter that are also part of hPINs filtered with increasingly stringent disparity thresholds. Fig. 5a reports that for significance levels up to α ≈ 0.38, the disparity-filtered network includes almost all edges with weights ≥ 2 (*w*_*c*_ = 2). For significance levels close to 0, the resulting hPIN contains almost all interactions with weights ≥ 6. This means that the disparity filter retains protein interactions supported by many pieces of evidence, which are more likely to be true positives^15^. Note that Fig. 5a emphasises the main difference between global and disparity filtering: while the former eliminates all interactions with weights below *w*_*c*_, the latter removes only those whose weights are not significant according to α. Because of this, we can have a disparity-filtered hPIN with, for example, some PPIs of weight 2 and the rest of at least 3 (see Fig. 5a). This is not possible in networks pruned with the global filter, which either contain all interactions of weight 2 or none of them.

**Figure 2.**
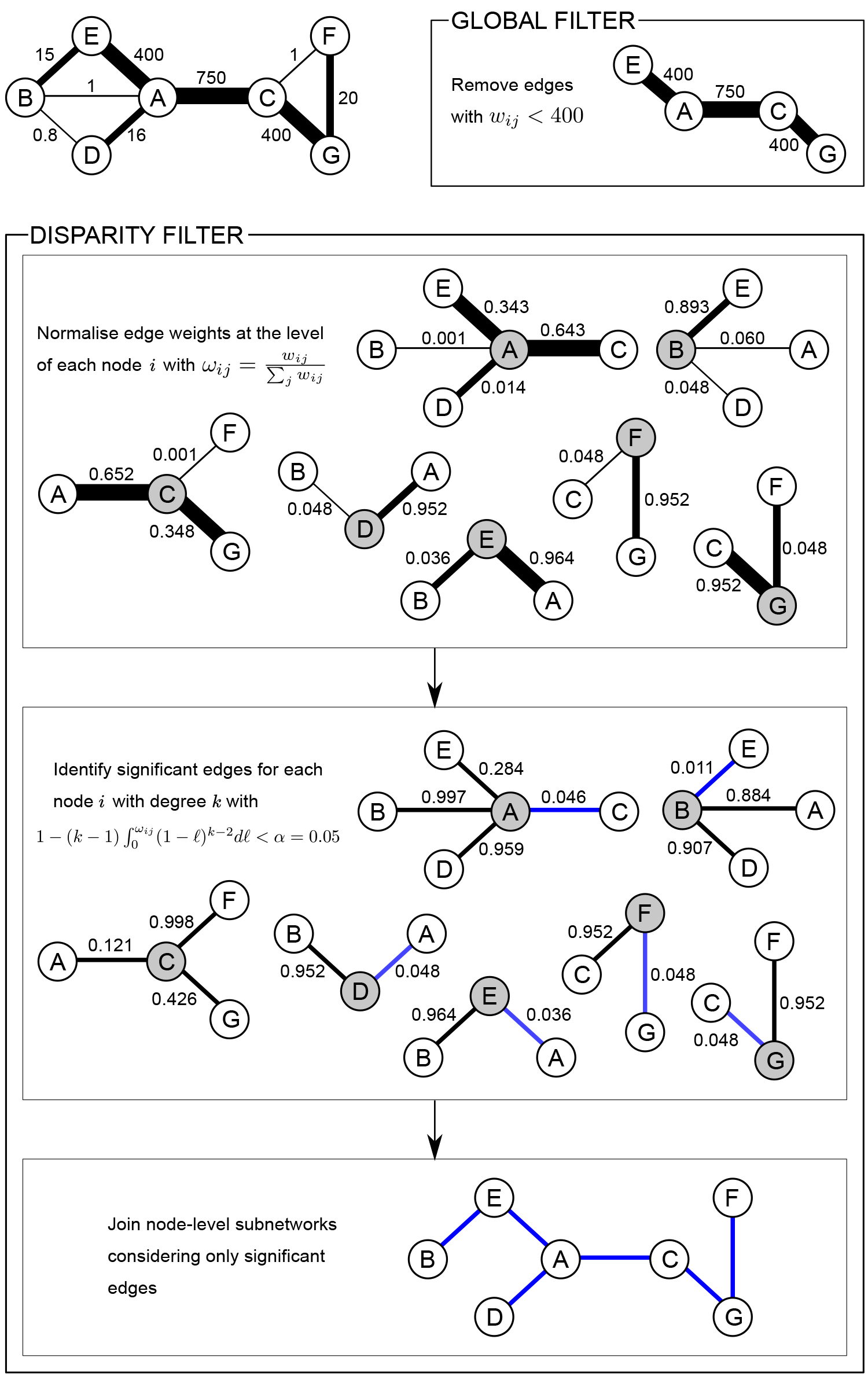
Global and disparity filtering. Illustrative example of the application of the global and disparity filters to an undirected weighted network. Using the upper quartile of the weight distribution (400) to remove edges with small weights results in the loss of nodes B, D and F and a network connected by the heaviest edges only. In contrast, applying the disparity filter with a significance level α = 0.05 results in a network that retains all nodes and the most significant edges (highlighted in blue).

**Figure 3.**
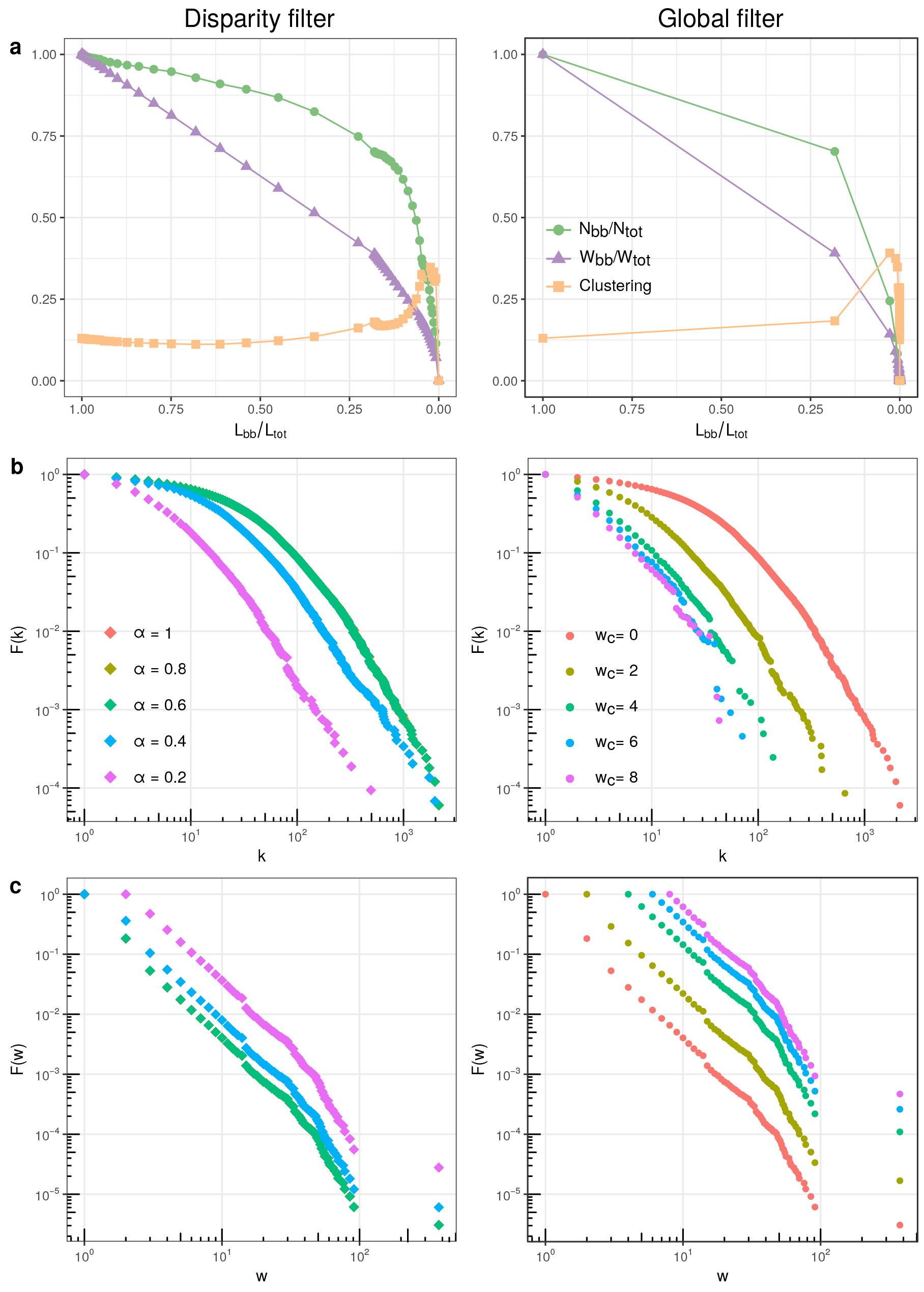
Global and disparity filters applied to the hPIN. (a) The remaining fraction of proteins (*N*_*bb*_/*N*_*tot*_), the remaining fraction of total weight (*W*_*bb*_/*W*_*tot*_) and the network’s clustering coefficient as a function of the remaining fraction of protein interactions (*L*_*bb*_/*L*_*tot*_) resulting from the application of different stringency levels of the disparity and global filters to the hPIN. (b) Changes in the cumulative degree distribution as different disparity and global thresholds are applied to the hPIN. (c) Same as **b** but for the cumulative weight distribution.

**Figure 4.**
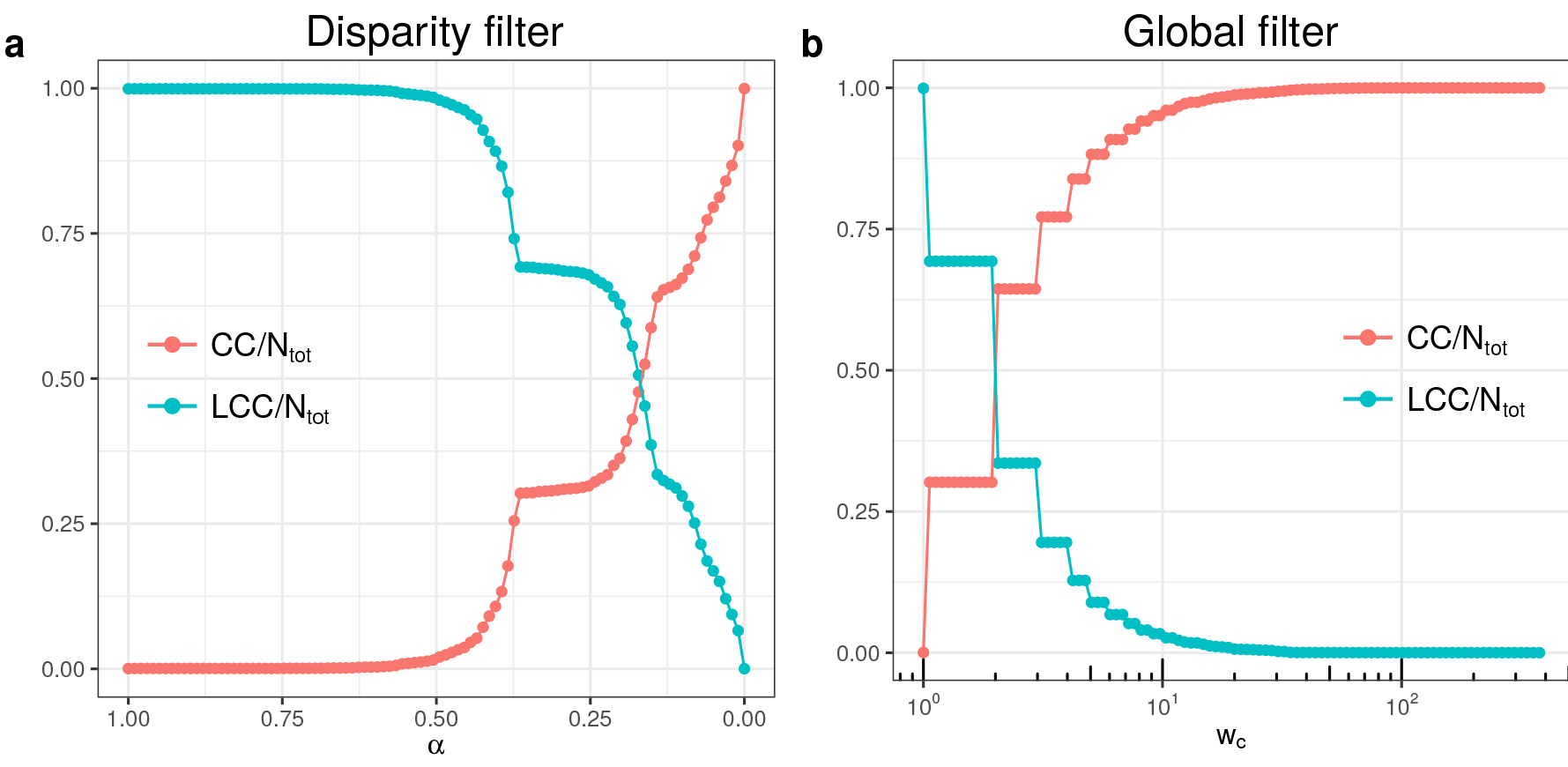
Connected components of filtered hPINs. (a) Changes in the number of connected components (*CC*) and size of the largest connected component (*LCC*) relative to the total number of proteins (*N*_*tot*_) as more stringent disparity thresholds α are applied to the hPIN. (b) Same as **a** but for increasing global thresholds *w*_*c*_.

We also contrasted the values of α for each edge in the hPIN with its HIPPIE and STRING confidence scores (see Methods). For this, we binned the significance values in ten equally sized groups and, as shown in Fig. 5b, it is the lowest αs that are related to the highest confidence scores. We can then conclude that the disparity-filtered network is of high quality and contains the most relevant and reproducible information.

### Bias reduction with the disparity filter

As mentioned in the Introduction and shown in Fig. 1, there is a clear correlation between the degree of a protein in the hPIN and how well-studied it is. This study bias has grave consequences on the different areas where network biology is playing an increasing role. Consider, for example, the network-based identification of disease-associated proteins^3–5,37^. A protein of interest is likely to interact with highly-connected entities in the hPIN that are, say, cancer-related. Nevertheless, the association of our protein with cancer might just be an effect of the network’s study bias. If we want to avoid incorrect conclusions stemming from this issue, it is thus imperative to unbias the human protein network.

In Fig. 6a we show how the correlation between protein degree and number of studies, as well as the slope of the linear fit shown in Fig. 1, decreases as we apply more stringent disparity filters α to the hPIN. The values of correlation and slope reach a minimum at α_*c*_ ≈ 0.38 and after this significance level, they increase again, probably due to random contributions from the few remaining proteins in the network^33^. We used α_*c*_ to disparity-filter the hPIN and retain proteins with a more homogeneous number of associated studies irrespective of their degree. Fig. 5 shows that this significance level retains the most reliable PPIs.

It is important to note that when we filter the hPIN using α_*c*_, isolated proteins (i.e. proteins with degrees *k* = 0) are removed from the network. Fig. 6b shows that discarded nodes represent the poorly-studied proteins from the original topology. Moreover, these proteins are involved in biological processes, like transport and metabolism, that hint at their participation in very transient interactions that may be mediated by molecules other than proteins (see Supplementary Data S4).

The application of a global filter results in similar trends (see Fig. 6c,d). However, the *w*_*c*_ at which the minimum correlation occurs produces a network with only 3% of the original number of proteins (see Fig. 6c). The backbone extracted with α_*c*_, in contrast, is formed by 81% of the original nodes (*N*_*bb*_ = 13,480, see Fig. 6a), 31% of the original links (*L*_*bb*_ = 102,749) and retains 49% of the original network’s weight (see Supplementary Data S3). As of 16th October 2017, SwissProt, the manually curated protein knowledgebase, reports 20,237 human proteins^38^. This means that our backbone covers 67% of the human proteome and is two times bigger than most of the recently published large-scale hPINs ^13,17^.

Altogether, these results underscore how the disparity filter can correct for the study bias present in the hPIN, retaining the most significant and reliable protein interactions and discarding poorly-studied system components, which most likely add spurious interactions to the network structure.

**Figure 5.**
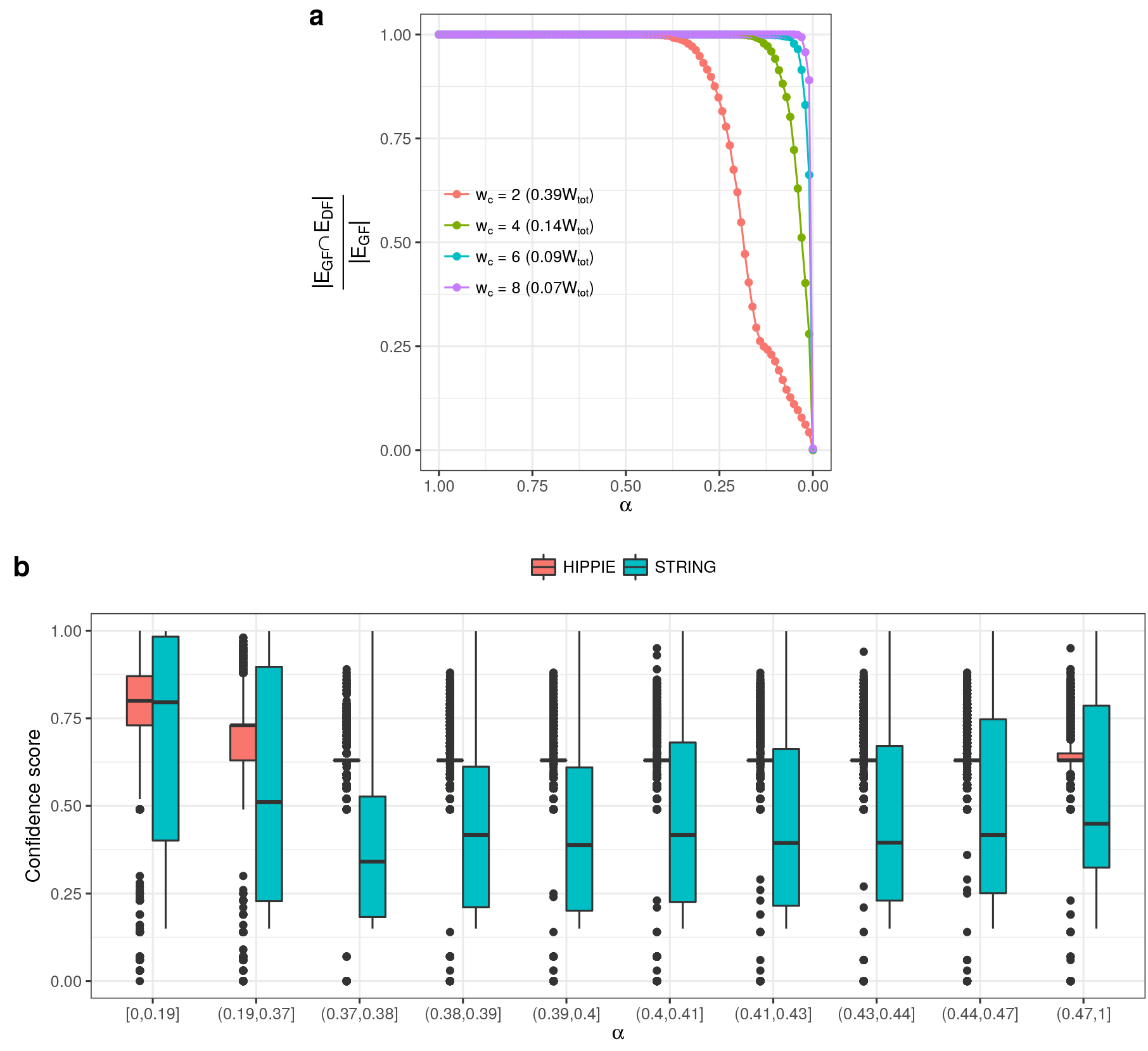
The disparity filter retains reliable protein interactions. (a) Fraction of protein interactions in four different hPINs pruned with the global filter (*E*_*GF*_) that are also present in hPINs filtered with increasingly stringent disparity thresholds α (*E*_*DF*_). The fraction of the original network weight (*W*_*t*_*ot*) that remains in the hPINs pruned with the global filter is reported in parentheses. (b) Salient protein interactions according to the disparity filter (i.e. edges whose weights yield low values of α) correspond to protein pairs with high confidence scores in the HIPPIE and STRING databases.

**Figure 6.**
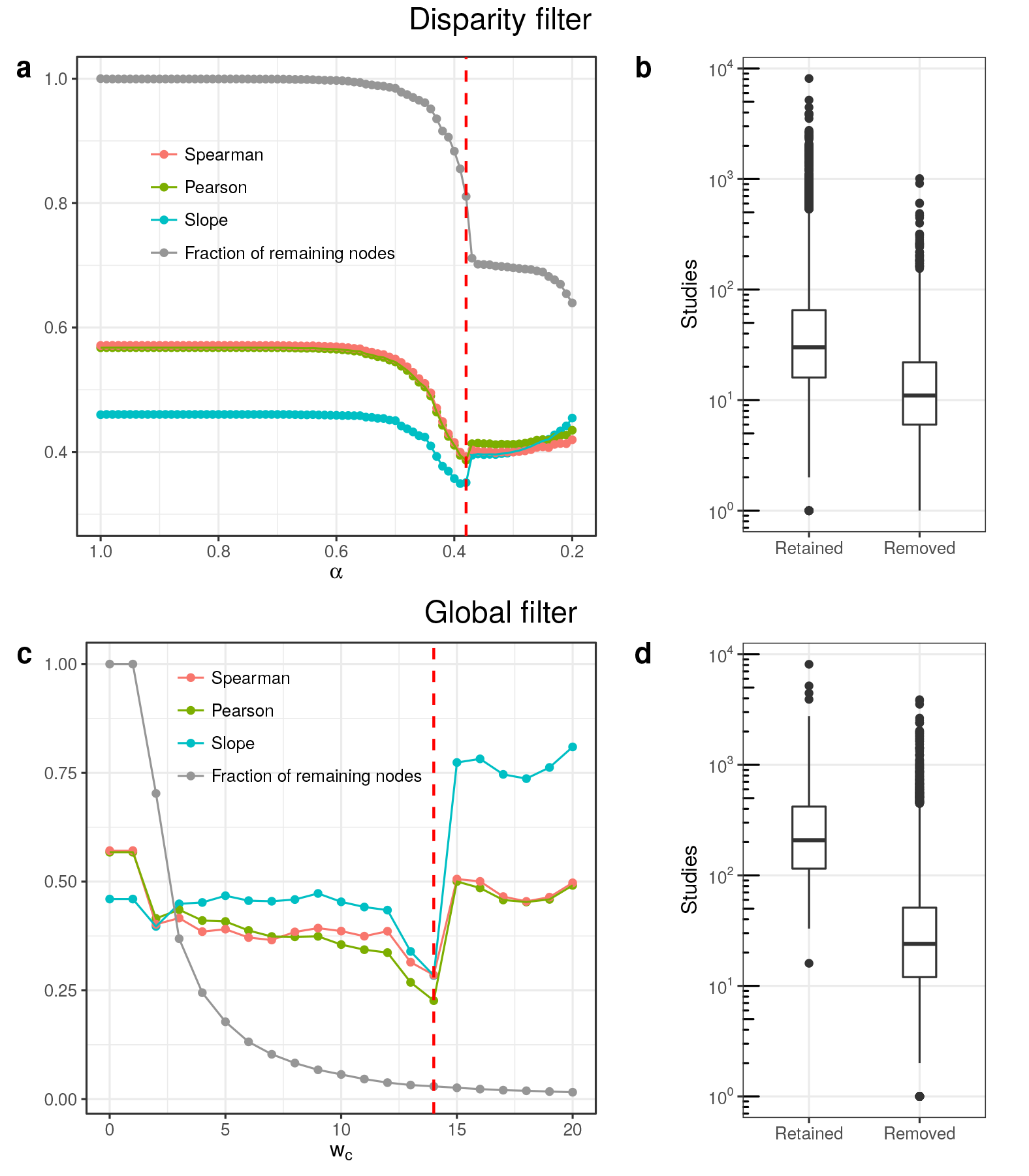
Bias reduction of the hPIN with the disparity filter. (a) As more stringent disparity thresholds α are applied to the hPIN, the Pearson and Spearman correlations between degree and number of studies are greatly reduced, as well as the slope of the linear fit shown in Fig. 1. The fraction of remaining nodes, relative to the original network, remains high. The red dashed line indicates the value of the threshold α_*c*_ that yields the lowest correlation between degree and number of studies. (b) Number of studies associated with retained and discarded proteins after the application of the disparity filter with α = α_*c*_. (c) Same as a but for a global filter that removes all protein interactions supported by less than *w*_*c*_ studies. (d) Same as **b** but for the global filter.

**Figure 7.**
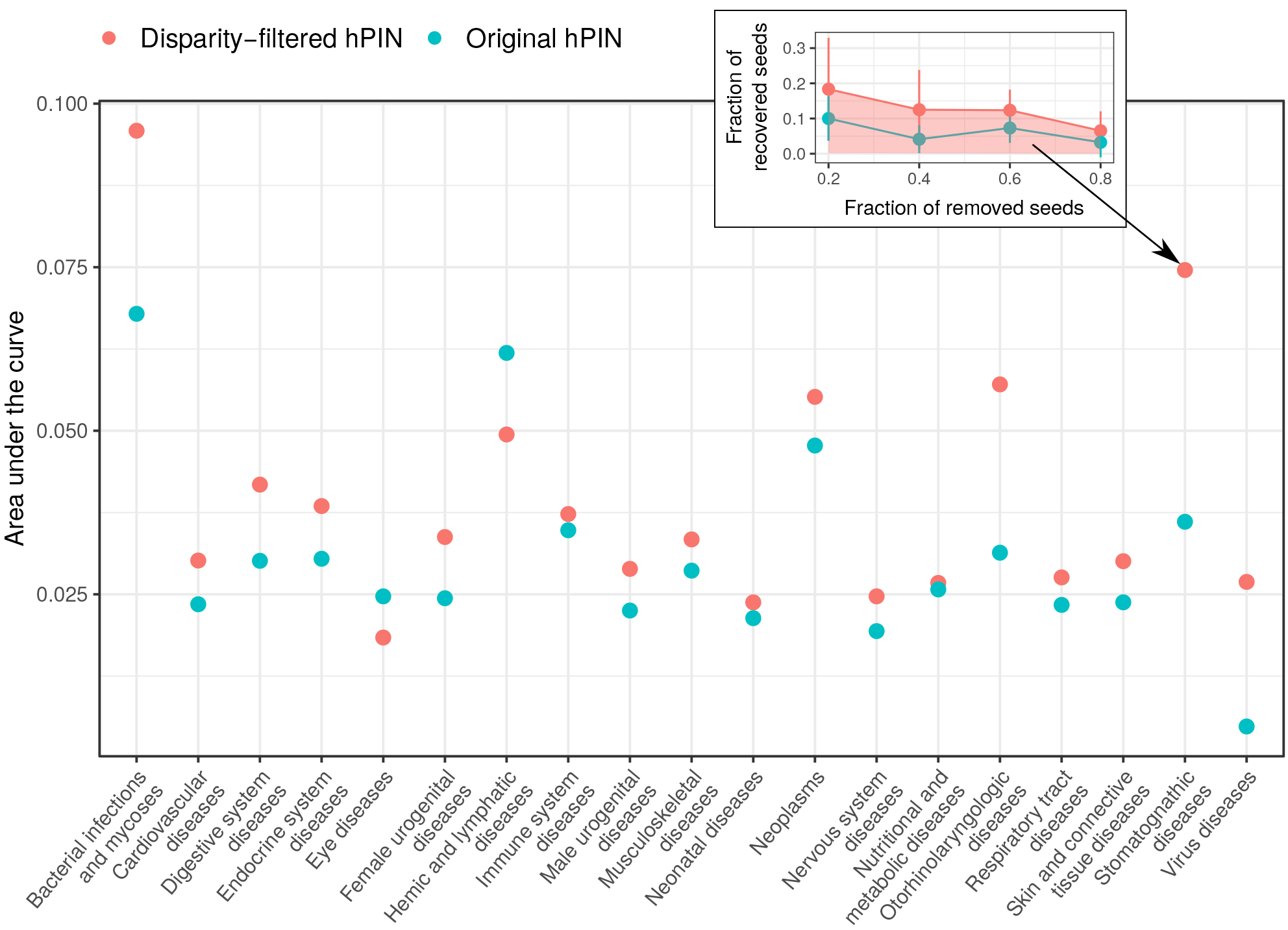
Growing disease modules with the filtered hPIN. An increasing fraction of proteins associated to 19 disease modules (seed proteins) was randomly removed from each one to later apply an algorithm that uses network connectivity patterns to identify potential disease module members. The algorithm was applied to the disparity-filtered network and to the unfiltered one. The inset shows the average fraction of recovered seeds associated to *Stomatognathic diseases* as a function of the fractions removed. The area under the curve for the disparity-filtered hPIN is highlighted. Error bars correspond to standard deviations. The plot reports areas under the curve for the rest of the disorders.

### Disease module detection with the unbiased hPIN

The emerging field of network medicine highly depends on a reliable hPIN for the study of how perturbations of proteins and their interactions may result in a disease phenotype. Recently, a myriad of methods has been developed to prioritise potential disease proteins based on their connectivity features in the hPIN^3–5,37^. These approaches rely on the disease module hypothesis, which states that gene products involved in a particular disease phenotype segregate in the same neighbourhood of the hPIN^8,39^. As a result, the application of these algorithms to noisy and biased networks can derive in spurious protein-disease associations.

We investigated the differences between performing disease module detection on the unfiltered hPIN and on its corresponding disparity-filtered version (see the previous section). Our analysis revolved on the application of the DiseAse MOdule Detection (DIAMOnD) algorithm^5^ to 19 disease modules (DMs) compiled by Menche and colleagues^39^ (see Methods and Supplementary Data S5). Given a list of known disease-related proteins (seeds) and a protein interactome, DIAMOnD grows the disease module with gene products that are significantly connected to it ^5^.

We randomly removed an increasing fraction of seeds from each disease and retrieved 200 candidate module members from the original and disparity-filtered hPIN using DIAMOnD. From these 200 candidates, we computed the fraction that coincides with the set of removed seeds and repeated the experiment 10 times (see inset in Fig. 7). Fig. 7, shows the area under the described curve and reports that in 17 out of the 19 analysed cases (89%), it is better to grow disease modules with the disparity-filtered network. This result stresses the value of the proposed filtering method in a field where a high-quality hPIN structure is crucial.

## Conclusions

A reliable, unbiased proteome-scale human protein network will play a pivotal role in pushing the boundaries of what we currently know about the living cell. Projects to generate such a reference hPIN are already in place^15,17^ and testing for the interaction of all possible protein pairs may be completed in the next few decades^13^. In the meantime, efforts like STRING^19^, HIPPIE^21^ and the proposed disparity-filtered hPIN can pave the way for the future of systems and network biology.

The present work resorts to the disparity filter to not only identify salient protein interactions but also produce a less-biased hPIN structure. Contrary to the excessively pruned topologies resulting from the removal of connections with weights below a given cutoff, the disparity filter identifies interactions with weights representing a significant fraction of each protein’s local strength. The extracted backbone contains most of the human proteins with reported PPIs, decouples node degree from number of studies and retains high-quality protein interactions according to reference databases. Moreover, we demonstrated how disease module detection methods can improve their precision with the disparity-filtered hPIN and anticipate that other areas of systems biology will benefit from this method to increase the quality of the hPIN.

## Methods

### The null model of the disparity filter

As stated in the Results, salient links in the disparity filter are those whose normalised weight is significantly greater than a distribution of randomly generated weights. For a node of degree *k*, these random weights are the result of distributing *k* − 1 points over the interval [0,1] uniformly at random^33^. The length of the *k* segments into which [0,1] is divided is thus distributed according to *p*(*ℓ*) = (*k* − *1*)(*1* − *ℓ*)^*k*−2^. This expression can be derived from a more general problem definition.

For *a* ∈ ℝ_>0_ and *k* ∈ ℕ_>0_ draw *k* − 1 points *X*_1_, …,*X*_*k*−1_ independently and uniformly at random from the interval *I* = [0, *a*]. These *X*_*i*_ partition *I* into *k* segments of lenght *Y*_*j*_, such that 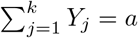. We are interested in how these *Y*_*j*_ are distributed.

Since *X*_*i*_ are uniformly and independetly placed over [0, *a*], they have the same density and cumulative distribution functions: 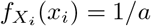 and 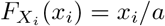 for *x*_*i*_ ∈ [0, *a*], respectively.

Let us focus on the rightmost segment *Y*_*k*_, whose length is *a* − max(*X*_*i*_). The probability that *Y*_*k*_ is larger than *ℓ* is:

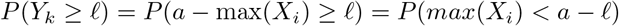

Since all *X*_*i*_ have the same cumulative distribution function, this last expression is equivalent to:

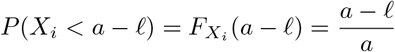

Thus, the joint distribution for the *k* − 1 independent *X*_*i*_ is 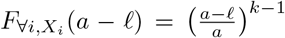 and we can now compute:

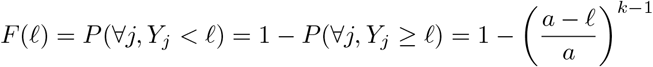

After differentiation, we obtain the probability density function:

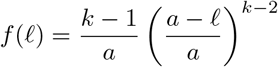

For the disparity filter, we partition the interval [0,1], i.e. *a* = 1, and *p*(*ℓ*) follows.

### Protein interaction network construction

The network used here consists of all experimentally-supported interactions between human proteins reported by PSICQUIC interaction providers^36^ as of 18th July 2017. PSICQUIC is an effort of the Human Proteome Organisation-Proteomics Standard Initiative (HUPO-PSI) to facilitate programmatic access (URLs and scripts) to molecular interaction databases. Using this programming interface, we computed the number of studies supporting each protein interaction and took it as the edge weight in the constructed human protein network. In addition, we retrieved each interaction’s confidence score from release 2.1 of the Human Integrated Protein-Protein Interaction rEference (HIPPIE)^21^ and from release 10.5 of the STRING database^18^.

### Studies associated with each protein

The studies associated with a protein denote the total number of PubMed identifiers associated with its Entrez Gene ID. This information is available in the NCBI Gene database^40^ at ftp://ftp.ncbi.nlm.nih.gov/gene/ DATA/gene2pubmed.gz. Our results correspond to the 4th May 2017 snapshot of gene2pubmed.

### Gene Ontology and Reactome pathway enrichment analyses

We carried out Gene Ontology^41^ and Reactome pathway^42^ enrichment analyses with R package FunEnrich, which is available at https://github.com/galanisl/FunEnrich.

### Protein-disease associations

We used a reduced version of the protein-disease associations compiled by Menche and colleagues^39^. The original dataset reports disease modules (groups of proteins associated with a condition) for very specific human disorders that, in some cases, are formed by only a few proteins. In consequence, we reduced it to 19 disease categories by merging modules that are perfect subsets of larger ones.

### The DIAMOnD algorithm

DIAMOnD detects novel disease-related proteins by identifying unexpected connectivity patterns between gene products and a disease module of interest. To this end, given a protein interactome and a set of known disease-related proteins (seeds), DIAMOnD focuses on gene products with direct links to the disease module. Then, via a Fisher’s exact test, it determines which of these proteins has the most significant number of connections to the module and adds it to the set of seeds. This process can be repeated several times to expand the number of seeds as desired. We applied DIAMOnD with 200 iterations. For more details see^5^.

### Hardware used for experiments

We executed all the experiments presented in this paper on a Lenovo ThinkPad 64-bit with 7.7 GB of RAM and an Intel Core i7–4600U CPU@2.10 GHz ⨯ 4, running Ubuntu 16.04 LTS. The only exceptions were the disease module detection experiments, which were executed on nodes with 64 GB of RAM, within the Mogon computer cluster of the Johannes Gutenberg Universität.

### Code availability

An R implementation of the disparity filter is available at https://github.com/galanisl/DisparityFilter. A python implementation of the DIAMOnD algorithm is available at https://github.com/barabasilab/DIAMOnD.

### Data availability

The following data accompanies this text:

- **Supplementary Data S1.** List of proteins in the human protein interaction network (hPIN) and their number of associated studies.
- **Supplementary Data S2.** The hPIN.
- **Supplementary Data S3.** The hPIN after the application of the disparity filter with α_*c*_.
- **Supplementary Data S4.** List of proteins removed from the hPIN after the application of the disparity filter, together with their Gene Ontology and pathway enrichment analysis.
- **Supplementary Data S5.** Protein-disease associations.

## Acknowledgements

We gratefully acknowledge the computing time granted on the supercomputer Mogon at the Johannes Gutenberg Universität. We also thank Prof. M. Ángeles Serrano for her valuable comments and suggestions.

### Competing interests

The authors declare that they have no competing interests.

### Author’s contributions

GAL integrated all the data, implemented and carried out the experiments. MAAN supervised the research. GAL wrote the manuscript, incorporating comments, contributions and corrections from MAAN. All authors read and approved the final version of the manuscript.

